# Human papillomavirus type 16 circular RNA is barely detectable for the claimed transformation activity

**DOI:** 10.1101/2021.09.22.449294

**Authors:** Lulu Yu, Zhi-Ming Zheng

## Abstract

The human papillomavirus type 16 (HPV16) E7 oncoprotein plays an essential role in cervical carcinogenesis and is encoded predominantly by a viral E6*I mRNA through alternative RNA splicing of a p97 promoter-transcribed bicistronic E6E7 pre-mRNA. Recently, Zhao et al. detected the HPV16 circular RNA circE7, which is generated aberrantly through backsplicing of the E6E7 pre-mRNA from two HPV16-positive cervical cancer cell lines, CaSki and SiHa. Based on their findings that HPV16 E7 was translated from circE7 and knockdown of circE7 in CaSki cells led to reduction of E7 oncoprotein, cell proliferation, and xenograft tumor formation, the authors claimed that circE7 is functionally important in cell transformation. We believe, however, that the reported circE7 function is overstated. We found that circE7 in CaSki cells is only 0.4 copy per cell and determine that the claimed circE7 function in the published report was resulted from off-targeting viral E7 linear mRNAs by their circE7 small interfering RNAs.

---

High-risk HPVs utilize a major early promoter to transcribe early transcripts for translation of different early proteins by alternative RNA splicing^1,2^. The viral E6 and E7 oncoproteins are responsible for disrupting cell cycle control and promoting cell proliferation and transformation activities of high-risk HPVs through interactions with multiple cellular factors^3,4^ and regulation of expression of non-coding RNAs^5,6^. However, production of E6 and E7 proteins is dependent on alternative splicing of the bicistronic E6E7 pre-mRNAs, and this RNA splicing is extremely efficient in HPV16- and HPV18-positive cervical cancer cells^7-10^. As shown in Fig.1a, HPV16 E6 and E7 are transcribed from the same early promoter P97 as a single bicistronic E6E7 transcript containing three exons and two introns, with intron 1 in the E6 open reading frame (ORF) bearing one splice donor (SD) at nucleotide (nt) 226 and two alternative splice acceptor (SA) sites, one at nt 409 and the other at nt 526, and intron 2 in the E1 ORF starting from an SD site at nt 880 in the HPV16 genome. Only the intron 1-retained E6E7 RNA has an intact E6 ORF and thus is capable of translating full-length E6 protein, but this RNA translates no or very little E7 due to intercistronic space restriction, of which the E6 stop codon is separated from the E7 start codon by only 2 nt. This limited intercistronic space prevents the translating ribosome from re-initiating translation of the downstream E7 ORF after stopping E6 translation. We have previously demonstrated that the HPV16 major spliced isoform E6*I RNA functions as E7 mRNA for production of E7 protein because the intron 1 splicing from nt 226 to nt 409 introduces a premature stop codon downstream of the splice junction, which thereby increases the intercistronic distance to 145 nt between the E6*I stop codon UAA and the E7 initiation codon AUG^7-9^. This phenomenon also applies to HPV18 E7 production^10^.

All introns in a pre-mRNA are spliced in 5’ SD to 3’ SA order as an intermediate circular lariat RNA^9,10^. Occasionally, RNA backsplicing from a downstream intron 5’ SD to an upstream intron 3’ SA over the exon(s) could occur, producing circRNAs containing the exon(s)^11^. Efficiency of RNA backsplicing to produce circRNAs is very low and may account for <1% of normal linear RNA splicing^12,13^. In the original article, Zhao et al. detected circE7 from the backsplicing of HPV16 E6E7 mRNAs in CaSki and SiHa cells and in cervical tissue samples (Fig.1a, lower right). The authors found that circE7 could translate E7 protein and knocking down circE7 expression by short hairpin RNAs (shRNAs) targeting to the backsplice junction in CaSki cells led to reduction of E7 protein expression and subsequent cell proliferation or xenograft tumor formation. However, this article is hard to follow because of mislabeled figure panels (Supplementary Figs. 2b, 2c, 4a, 5a, and 5b), a missing figure legend (Supplementary Fig. 5f), no size markers for RT-PCR products (Figs. 1d, 2b, 2e, 3g, 5d, and 5e and Supplementary Figs. 2c, 3c, and 3g) and Northern blots (Supplementary Figs. 2b and 4b), misuse of the circE7 g-block sequence which had four, including E7, AUG mutations as a wild-type circE7 (supplementary data 2), and lack of suitable controls. In addition, the authors utilized unconventional HPV16 genome positions to describe their circE7 production and viral RNA splicing. We had to align their HPV16 genome positions to an HPV16 reference genome from the papillomavirus database PAVE (http://pave.niaid.nih.gov) in order to understand how their studies were designed and performed.

**Fig. 1.**
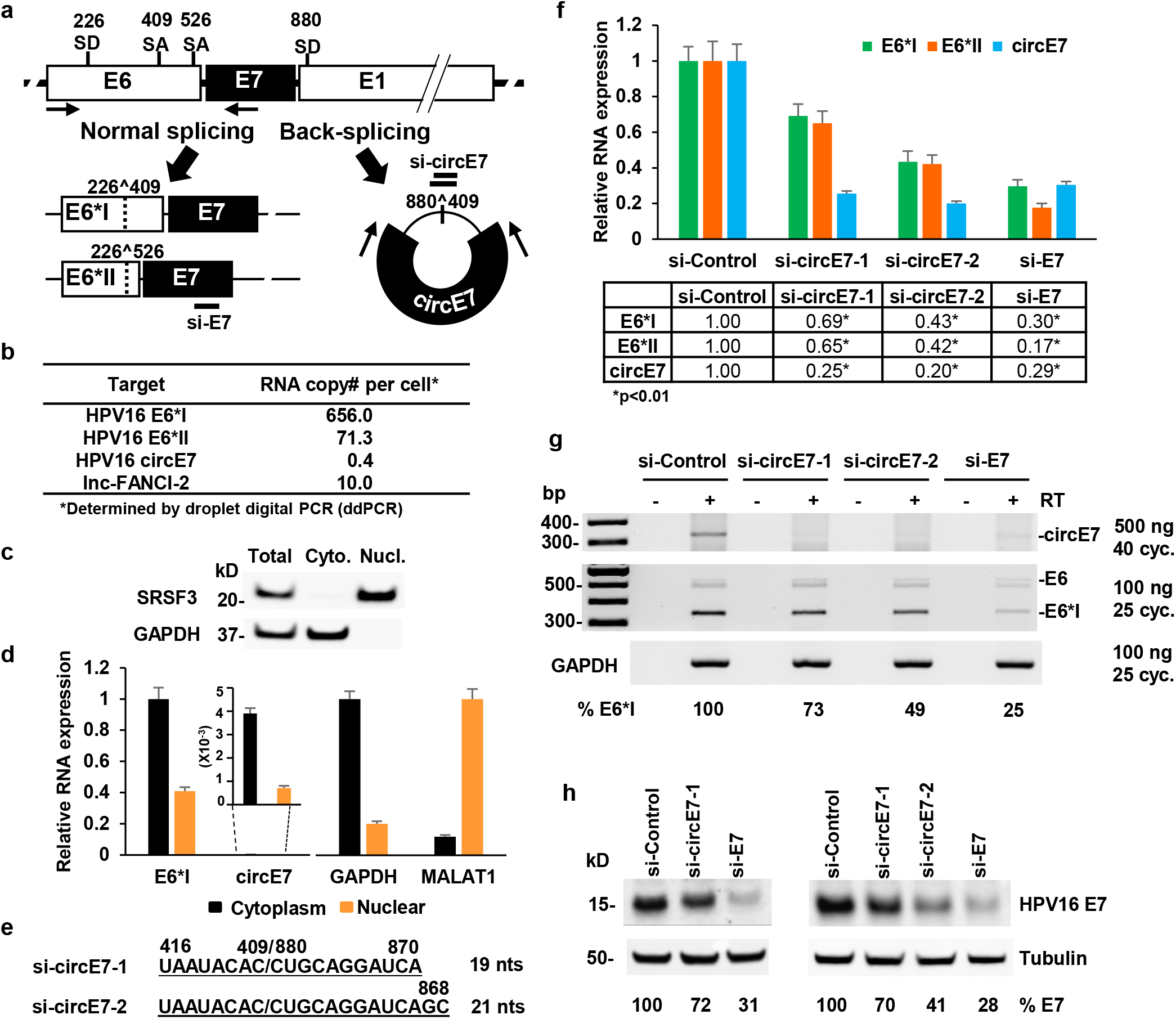
Characterization of HPV16 circE7 expression in CaSki cells. **a** Structure diagram of viral early promoter p97-derived bicistronic HPV16 E6E7 pre-mRNA and its splicing profiles in CaSki cells. Shown on the pre-mRNA are E6, E7, and partial E1 ORF, and genomic positions of viral splice donor (SD) and splice acceptor (SA) sites. Below the pre-mRNA are two spliced isoforms (E6*I and E6*II) from alternative RNA splicing commonly detected in CaSki cells and a barely detectable circE7 from backsplicing. Paired arrows below the pre-mRNA and on circE7 RNA were used for detection of individual products from viral RNA splicing in CaSki cells. Horizontal bars above the backsplice junctions and below the E7 ORF are siRNAs used to target circE7 (si-circE7-1 and −2) and the corresponding RNA isoforms containing the E7 ORF region in this study. **b** RNA copy number of HPV16 E7 (E6*I and E6*II) and circE7 in CaSki cells determined by ddPCR. Total cell RNA was reverse transcribed and serially diluted in triplicate before ddPCR analysis, with long noncoding RNA lnc-FANCI-2 serving as the internal reference RNA control. **c** and **d** HPV16 E7 (E6*I) and circE7 RNAs in CaSki cells are predominantly cytoplasmic. Nuclear and cytoplasmic fractionation efficiency was determined by Western blot analysis of nuclear protein SRSF3 and cytoplasmic GAPDH protein (**c**) and by RT-qPCR analysis of nuclear MALAT1 and cytoplasmic GAPDH RNA (**d**). E7 (E6*I) and circE7 RNAs in nuclear and cytoplasmic fractions were determined by RT-qPCR (**d**). Data are mean + SE of six repeats. **e** Nucleotide sequences of two synthetic siRNAs targeting the circE7 backsplice junction. **f, g**, and **h** HPV16 circE7-specific siRNAs affect the expression of both E6*I and E6*II RNAs and E7 protein in CaSki cells. Cell lysates were prepared 72 h after transfection of indicated individual siRNAs. Total RNA extracted from the cells was reverse transcribed and used to quantify HPV16 E6*I, E6*II, and circE7 RNAs by RT-qPCR using specific TaqMan probes (**f**). Data are mean + SE of six repeats. *, p<0.01 by unpaired, two-tailed Student’s *t* test. The cDNAs prepared were also used to profile the effects of individual siRNAs on viral circE7 and bicistronic E6E7 RNA splicing by RT-PCR and agarose gel electrophoresis. A total of 500 ng cDNA and 40 cycles of PCR reactions were used to detect circE7, and 100 ng of cDNA and 25 cycles of PCR reactions were used to detect the common spliced products of E6E7 and cellular GAPDH pre-mRNAs. GAPDH served as the RNA loading control. The relative E6*I RNA level was quantified after normalizing to GAPDH in each sample (**g**). Total cell lysates were also immunoblotted for E7 protein expression by an anti-HPV16 E7 antibody. The relative E7 level in each sample was quantified after normalizing with β-tubulin serving as the protein loading control (**h**).

We have been studying the RNA splicing of both HPV16 and HPV18 E6E7 pre-mRNAs^7-10,14-16^. We recently quantified by ddPCR the copy number of HPV16 E6*I, E6*II, and circE7 in CaSki cells using human lnc-FANCI-2 with a known RNA copy number per CaSki cell^6^ as a reference RNA and found only 0.4 copy of circE7 per CaSki cell, which is ∼1640-fold less than E6*I mRNAs (Fig. 1b). The number of E6*I transcripts in CaSki cells is ∼10 times higher than E6*II as we reported^15^. Although the circE7 RNA produced from backsplicing is 472 nt long, the detected circE7 products from RT-PCR with an inverse primer pair used in our study was about 351 nt and appeared not to be enriched by RNase R treatment (Supplementary Fig. 1a). Approximate 500 ng of total cDNA and 40 cycles of inverse PCR were applied to detect circE7 because of its extremely low abundance in CaSki cells (Supplementary Fig. 1a) and two W12 subclone cell lines (Supplementary Fig. 1b), when compared for detection of HPV16 E6*I and E6*II RNAs (Supplementary Fig. 1a). The RT-PCR products were confirmed as circE7 by Sanger sequencing. Although Zhao et al. claimed detection of circE7 by Northern blot analysis using an RT-PCR product as a probe, lack of further verification of the reported Northern blot band (∼700 nt), larger than its predicted size, indicated that this entity might not be authentic.

We further showed that the detected E6*I and circE7 in CaSki cells are both cytoplasmic by cell fractionation analysis (Fig. 1c-d). As expected, the level of circE7 RNA in the cytoplasm is negligible when compared to E6*I (Fig.1d). The majority of HPV16 early transcripts and spliced RNA isoforms have been identified in the cytoplasm by cell fractionation and Northern blot analysis^14^. Surprisingly, Zhao et al. found viral linear E7 RNA was detected predominantly in the nuclear fraction by SYBR green RT-qPCR both in CaSki cells and in E6E7 minigene transfected HEK293T cells (Zhao et al., Fig. 3b). We analyzed their primer sequences and found that the primer pair detected only less abundant intron 1-containing E6 RNA or most likely the nuclear E6E7 pre-mRNAs, but not the predominant E6*I RNAs in CaSki cells. It is also hard to determine the exact amount of linear E7 RNA that was expressed from each minigene in HEK293 cells by SYBR green RT-qPCR, as everything in the report was shown as a ratio and only the circE7 RNA was displayed by inverse PCR and in Northern blots (Zhao et al., Fig. 3a). Unless a complete expression profile is provided, it will be difficult to verify that the detected E7 protein was only translated from circE7, not from linear E7. In fact, addition of the QKI binding site to facilitate the circulation of E7 RNA only enhanced production of circE7 by two-fold (Zhao et al., Supplementary Fig. 3c). We also question why two siRNAs targeting to the linear E7 RNA expressed from the minigenes led to its reduction by >75% (Zhao et al., Supplementary Fig. 3e) but did not decrease the expression of FLAG-E7 protein (Zhao et al., Fig. 2c). Clearly, the authors imply that their linear E7 RNAs somehow did not translate E7 protein. We found that the circE7 g-block sequence used in the authors’ study as a wild-type circE7 had four, including E7, AUG mutations (Zhao et al., Supplementary data 2).

We then investigated how circE7 RNA at 0.4 copy per CaSki cell could exhibit the high impact on E7 production and activities in cell proliferation and xenograft tumor formation reported by Zhao et al. We designed two siRNAs targeting the backsplice junction of HPV16 circE7 (Fig. 1e), with si-circE7-2 being the same circE7-sh2 sequence in the authors’ study. After delivery of the siRNAs to CaSki cells, we found both si-circE7-1 and si-circE7-2 could knock down circE7 expression by ∼75-80%, but si-circE7-1 also reduced E6*I and E6*II expression by 30-35% and si-circE7-2, which is 2 nt longer than si-circE7-1, decreased E6*I and E6*II expression up to 60%, whereas si-E7 targeting to the E7 ORF region^8^ inhibited ∼70% expression of E6*I, E6*II, and circE7 (Fig. 1f). We also found that all three siRNAs affected the expression of viral E6 RNAs by 15-20% (Supplementary Fig. 2). Splicing profile analysis by RT-PCR and agarose gel electrophoresis confirmed the siRNA knockdown effects on both circE7 and E6*I RNAs (Fig. 1g). Western blot analysis of the siRNA-treated CaSki cells further confirmed these three siRNA-mediated reductions of E7 protein expression in the order of si-E7 > si-circE7-2 > si-circE7-1 (Fig. 1h). The data are striking because we found that si-circE7-2 with 13-nt sequence overhang to the backsplice junction had a bigger off-target effect on linear E6*I and E6*II RNAs than si-circE7-1 with the 11-nt overhang sequence. Although Zhao et al. claimed that their shRNAs had no effects on linear E6/E7 or E6*I RNAs in CaSki cells, lack of an internal RNA loading control and probe information in the Northern blot (Zhao et al., Supplementary Fig. 4b) and miscalculation of circE7 shRNAs on E6*I expression by unusual normalization to β-actin in the SYBR green RT-qPCR (Zhao et al., Supplementary Fig. 4c) raised a red flag for their data interpretation. As the authors’ shRNA study can’t unequivocally prove E7 protein production only from circE7 at 0.4 copy per CaSki cell and their data conflict from our results, we believe that their claim of circE7 function in production of E7 protein and E7 activities was attributable to the off-target effects of their si-circE7s.

## Acknowledgements

This study was supported by the Intramural Research Program of the National Institutes of Health, National Cancer Institute, Center for Cancer Research. We thank Anne Arthur for proofreading and other members of the Zheng lab for comments and suggestions.

## Author contributions

L.Y. and Z.M.Z designed the study, performed all data analyses/interpretations, and wrote the manuscript.

## Competing interests

The authors declare no competing interests.

## Data availability statement

All the data in the manuscript is available upon request.

## Supplementary information

### Methods

#### Cell lines and siRNAs

HPV16-positive cervical cancer cell line CaSki was obtained from ATCC (Manassas, VA). CaSki cells were maintained in Dulbecco’s modified Eagle’s medium (DMEM; Thermo Fisher Scientific, Waltham, MA) with 10% fetal bovine serum (FBS; GE Healthcare, Logan, UT) at 37 °C in a 5% CO_2_ atmosphere. HPV16-positive W12 subclone cell lines 20861 (integrated HPV16 genome) and 20863 (episomal HPV16 genome) were a gift from Paul Lambert (University of Wisconsin). Individual subclones were maintained in F-medium (3:1, F-12:DMEM) supplemented with 5% FBS, 0.4 μg/ml hydrocortisone, 5 μg/ml insulin, 8.4 ng/ml cholera toxin, 10 ng/ml epidermal growth factor, 24 μg/ml adenine, 100 U/ml penicillin, and 100 μg/ml streptomycin. All subclone cells were grown in the presence of irradiated 3T3-J2 feeder cells. Three custom-designed synthetic siRNAs including two specifically targeting the backsplice junction of circE7 and one targeting the open reading frame of E7 (Supplementary Table 1) were designed and purchased from Dharmacon (Lafayette, CO). Non-targeting control siRNA (Dharmacon, #D-001210-01) served as the negative control. CaSki cells at 24 h of cell passage were transfected with 40 nM of siRNA by LipoJet In Vitro Transfection Kit (Ver. II, SignaGen Laboratories, Rockville, MD). Total protein extracts and total RNA were prepared 72 h after the siRNA transfection of CaSki cells. The cells were incubated with 10 µM MG-132 (#M7449, Sigma-Aldrich, St. Louis, MO) for 4 h before harvesting for protein samples.

#### RNA preparation and RNase R treatment

Total RNA from CaSki cells with or without siRNA treatment was extracted using TriPure reagent (#11667165001,Roche, Basel, Switzerland) according to the manufacturer’s protocol. Approximate 2 μg of RNA was treated for 40 min at 37 °C by RNase R (5 U) (#RNR07250, Lucigen, Middleton, WI) in 1× RNase R buffer.

#### Droplet digital PCR (ddPCR)

The number of HPV16 E6*I, E6*II, and circE7 RNA copies per cell was determined by ddPCR. Briefly, ∼2 μg of total RNA was reverse transcribed in a 20-μl reaction at 50 °C with random hexamers and SuperScript® IV Reverse Transcriptase (Thermo Fisher Scientific, #18091050). The human lnc-FANCI-2 RNA^6^ with known copy number per CaSki cell was used as a reference RNA to determine the cell number. The primers and TaqMan probes in ddPCR for detection of HPV16 E6*I, E6*II, circE7, and lnc-FANCI-2 are listed in Supplementary Table 1. The prepared CaSki cDNA was serially diluted at 1:2, 1:20, and 1:40 with a low-EDTA TE buffer (10 mM Tris-HCl, 0.1 mM EDTA, pH=8.0), and then 1 μl of each diluted cDNA was used in triplicate in a 20-μl ddPCR reaction containing the specific primer pair and a TaqMan probe for specific detection of HPV16 E6*I, E6*II, circE7, and lnc-FANCI-2. Droplet generation (Bio-Rad QX200) and PCR (Bio-Rad T100) were performed according to the manufacturer’s protocol (BioRad, Hercules, CA). The cycling protocol started with a 95 °C enzyme activation step for 10 min and followed by 40 cycles of a two-step cycling (94 °C 30 s and 60 °C 30 s at ramping rate of 2° C/s). The final extension time was 10 min at 98 °C. Bio-Rad QuantaSoft 1.5.38.1118 was used to process the data. The best dilution for circE7 detection was 1:2, while that for the other three targets was 1:40. The input cell number was determined by the level of lnc-FANCI-2. The mean HPV16 E6*I, E6*II, and circE7 copy number per cell was calculated from three ddPCR reactions in triplicate.

#### RT-PCR and RT-qPCR

Total RNA (2 μg) was reverse-transcribed as described above. PCR amplification was performed with Platinum™ SuperFi II DNA Polymerase (Thermo Fisher Scientific, #12361010) using the primers listed in Supplementary Table 1. 500 ng of cDNA was used for circE7 detection, and the cycling protocol was shown as follows: 98 °C 30 s, followed by 40 cycles of 98 °C 15 s, 55 °C 15 s, 72 °C 3 min, and the final elongation step of 72 °C 10 min. The same cDNA at 100 or 200 ng was used for linear E7 and GAPDH detection, of which the cycling protocol started with 98 °C 30 s, followed by 25 cycles of 98 °C 15 s, 55 °C 15 s, 72 °C 30 s, and the final elongation step of 72 °C 10 min. The following oligo primers were used in RT-PCR: oLLY405 and oZMZ212 for HPV16 circE7, oZMZ237 and oZMZ212 for HPV16 E6E7 and spliced RNA isoforms, and oZMZ269 and oZMZ270 for human GAPDH RNA. RT-qPCR was carried out using TaqMan Gene Expression Master Mix (Thermo Fisher Scientific, #4369016) together with the specific primers on a StepOne Plus Real-Time PCR system (Applied Biosystems, Foster City, CA). The customized RT-qPCR primers and probes for HPV16 E6*I, E6*II, and circE7 are listed in Supplementary Table 1. Human GADPH RNA (Thermo Fisher Scientific, #Hs02758991_g1) served as the RNA loading control.

#### CaSki cell fractionation

CaSki cells (5 × 10^6^) were fractionated by using Nuclei EZ Prep Kit (Sigma-Aldrich, #NUC-101) following the manufacturer’s protocol. Briefly, the cells in 10-mm dishes were rinsed with cold PBS (Thermo Fisher Scientific, #10010023), digested with 0.05% trypsin (Thermo Fisher Scientific, #25300062), quenched in DMEM medium containing 10% FBS, and then counted. The digested cells were then transferred to a 15-ml conical tube, centrifuged, and washed twice with cold PBS at 800 rpm for 10 min at 4° C. The cell pellet was transferred to a 1.5-ml Eppendorf tube and suspended in 600 μl of cold Nuclei EZ lysis buffer. Approximate 30 μl of the total cell lysate was collected for total cell protein detection. The remaining total cell lysate was centrifuged at 500 g for 5 min at 4 °C. The supernatant was then collected for cytoplasmic protein detection and RNA extraction using TriPure (Roche). The nuclei pellet was washed with 1 ml of Nuclei EZ lysis buffer and resuspended in 300 μl of Nuclei EZ storage buffer for nuclear protein detection and RNA extraction using TriPure (Roche). For Western blot analysis of fractionation efficiency, the fractionated cell lysates in 2× SDS loading buffer with 5% 2-ME were resolved by electrophoresis in a NuPAGE Bis-Tris 4–12% gel (Thermo Fisher Scientific) in 1× MOPS SDS buffer (Thermo Fisher Scientific) and immunoblotted for corresponding proteins by specific antibodies. The fractionated cytoplasmic and nuclear RNAs were used for detection of HPV16 E6E7 RNAs and circE7 by RT-qPCR with human GAPDH and MALAT1 (Thermo Fisher Scientific, #Hs00273907_s1) serving as controls for RNA fractionation efficiency of the cytoplasmic and nuclear RNAs.

#### Western blot analysis

Total cell protein for HPV16 E7 detection was prepared from 5 × 10^6^ CaSki cells by directly lysing the cells in 300 μl of 1× RIPA buffer (#BP-115X, Boston BioProducts, Ashland, MA) with the addition of cocktail protease inhibitors (chymotrypsin, thermolysin, papain, pronase, pancreatic extract, trypsin) (Roche, #04693159001). The cell lysate (15 μl) was mixed with 5 μl of 4× SDS loading buffer with addition of 5% 2-ME, heat denatured at 95° C for 10 min, and resolved by electrophoresis in a NuPAGE Bis-Tris 4–12% gel (Thermo Fisher Scientific) in 1× MOPS SDS buffer (Thermo Fisher Scientific). After transfer onto a nitrocellulose membrane, the membrane was blocked for 1 h with 5% skim milk in 1× TBS (Tris-buffered saline) containing 0.05% Tween (TTBS) and incubated with a primary antibody diluted in TTBS overnight at 4° C. After three washes with 1× TTBS buffer, the membrane was then incubated with a corresponding secondary peroxidase-conjugated antibody diluted in 2% milk/TTBS for 1 h at room temperature. The specific signal on the membrane was generated with SuperSignal West Pico (Thermo Fisher Scientific) and captured by a ChemiDoc Touch imaging system (Bio-Rad). To confirm the cytoplasm/nuclear separation in CaSki cells, CaSki cells in Nuclei EZ Lysis Buffer was directly lysed in 2× SDS loading buffer with addition of 5% 2-ME; all the other procedures are the same as shown above.

#### Antibodies used for Western blot analysis

Rabbit polyclonal anti-HPV16 E7 (#GTX133411, GeneTex, Irvine, CA), rabbit monoclonal anti-SRSF3 (#NBP2-76892, NOVUS, Littleton, CO), rabbit monoclonal anti-GAPDH (#14C10, Cell Signaling, Danvers, MD), and mouse monoclonal anti-β—tubulin (#T5201, Sigma-Aldrich) were used for Western blot analysis.

**Supplementary Table 1** DNA oligo primers, synthetic siRNA, and TaqMan probes used in the study.

---

**Supplementary Fig. 1.**
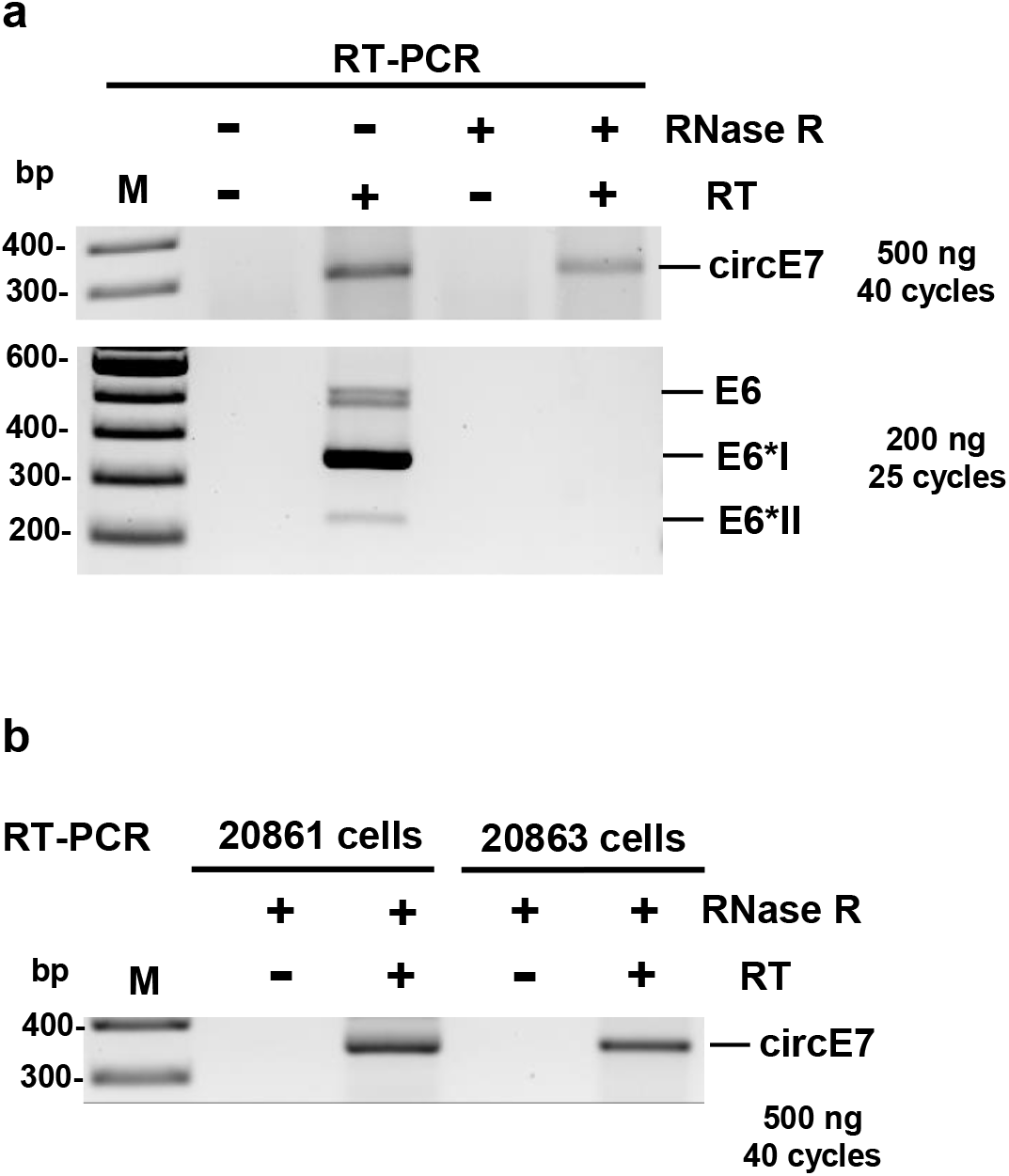
**a**. Detection of HPV16 circE7 and linear E7 in CaSki cells by RT-PCR upon RNase R treatment. RNase R (5 U)-treated total RNA was reverse transcribed in a 20 μl-reaction and 500 ng of the cDNA was then used for circE7 detection in 40 cycles of PCR amplification, whereas 200 ng of the cDNA was used for E6*I and E6*II detection in 25 cycles of PCR amplification. **b** Detection of HPV16 circE7 in W12 subclone 20861 cells with an integrated HPV16 genome and subclone 20863 cells with an episomal HPV16 genome by RT-PCR upon RNase R treatment. RNase R (5 U)-treated total RNA was reverse transcribed in a 20 μl-reaction and 500 ng of the cDNA was then used for circE7 detection in 40 cycles of PCR amplification.

**Supplementary Fig. 2.**
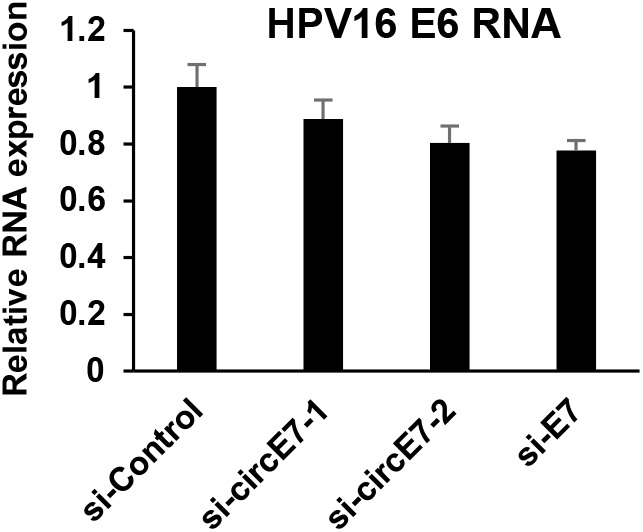
HPV16 circE7-specific siRNAs affect the expression of HPV16 E6 RNA in CaSki cells. Cell lysates were prepared 72 h after transfection of indicated siRNAs. Total RNA extracted from the cells was reverse transcribed and used to quantify HPV16 E6 RNA by RT-qPCR using an E6-specific TaqMan probe.

## Notes

### Competing Interest Statement

The authors have declared no competing interest.

